# Regulation of PP2A, PP4, and PP6 holoenzyme assembly by carboxyl-terminal methylation

**DOI:** 10.1101/2021.11.07.467570

**Authors:** Scott P. Lyons, Elora C. Greiner, Lauren E. Cressey, Mark E. Adamo, Arminja N. Kettenbach

## Abstract

The family of Phosphoprotein Phosphatases (PPPs) is responsible for most cellular serine and threonine dephosphorylation. PPPs achieve substrate specificity and selectivity by forming multimeric holoenzymes. PPP holoenzyme assembly is tightly controlled, and changes in the cellular repertoire of PPPs are linked to human disease, including cancer and neurodegeneration. For PP2A, PP4, and PP6, holoenzyme formation is in part regulated by carboxyl (C)-terminal methyl-esterification (often referred to as “methylation”). Here, we use mass spectrometry-based proteomics, methylation-ablating mutations, and genome editing to elucidate the role of C-terminal methylation on PP2A, PP4, and PP6 holoenzyme assembly. We find that the catalytic subunits of PP2A, PP4, and PP6 are frequently methylated in cancer cells and that deletion of the C-terminal leucine faithfully recapitulates loss of methylation. We observe that loss of PP2A methylation consistently reduced B55, B56, and B72 regulatory subunit binding in cancer and non-transformed cell lines. However, Striatin subunit binding is only affected in non-transformed cells. For PP4, we find that PP4R1 and PP4R3β bind in a methylation-dependent manner. Intriguingly, loss of methylation does not affect PP6 holoenzymes. Our analyses demonstrate in an unbiased, comprehensive, and isoform-specific manner the crucial regulatory function of endogenous PPP methylation in transformed and non-transformed cell lines

## Introduction

Dynamic protein phosphorylation is an essential regulatory mechanism that controls most cellular processes [1]. The opposing activities of protein kinases and phosphatases determine the phosphorylation state of proteins to control their activity and function. While serine and threonine phosphorylation is carried out by over 400 protein kinases [2], the majority of serine and threonine dephosphorylation is performed by the ten catalytic subunits that constitute the family of Phosphoprotein Phosphatases (PP1α, PP1β, PP1γ, PP2Acα, PP2Acβ, PP2Bc, PP4c, PP5c, PP6c, and PP7c) [3]. Many PPPs achieve substrate specificity through the formation of multimeric holoenzymes [1,4]. PP1α/β/γ, catalytic subunits bind to regulatory subunits to form heterodimers, while PP2Acα/β, PP4c, and PP6c form mostly heterotrimers with regulatory and scaffolding subunits. Holoenzyme formation is tightly regulated through various mechanisms, including post-translational modifications, to ensure the appropriate repertoire of PPPs is present in cells to catalyze specific dephosphorylation.

The catalytic subunits of PP2Acα, PP2Acβ, PP4c, and PP6c are closely related and highly conserved from yeast to human [3]. The PP2A trimer consists of a scaffold (A), regulatory (B), and catalytic (C) subunit [5]. Both the C and A subunits have two isoforms PP2Acα/β and PR65α/β, respectively. There is only an eight amino acid difference between the two catalytic subunit isoforms. Both isoforms bind A and B subunits equally well, and no difference in activity between them has been observed [6]. In cells and tissues, the A and B isoforms are ubiquitously expressed with the expression of the A isoform being several folds higher than the B isoform [7,8]. The A and C subunits of PP2A form a high-affinity dimer (AC) that binds to the regulatory B subunit. The regulatory subunit affects holoenzyme substrate specificity, cellular localization, and catalytic activity. There are four families of regulatory subunits: B55 (B/PR55), B56 (B’/PR61), B72 (B’’), and the striatin family (B’’’) [9,10]. Each family contains three to five different isoforms, and many isoforms have several splice variants. Combinatorially, PP2A subunits can form close to 100 unique holoenzyme trimers. The PP4 catalytic subunit (PP4c) forms heterodimeric and heterotrimeric complexes with five different regulatory subunits: PP4R1, PP4R2, PP4R3α, PP4R3β, and PP4R4 [11,12]. The PP6 catalytic subunit (PP6c) forms heterotrimeric complexes with one of three Sit4-associated protein subunits (SAPS1-3) and one of three Ankyrin repeat subunits (ANKRD28, ANKRD44, ANKRD52) [13,14].

Holoenzyme assembly is, in part, regulated by post-translational modifications (PTMs) of the catalytic subunits [15,16]. The C-terminus of PP2Acα/β, PP4c, and PP6c is highly conserved with the last three amino acids (tyrosine (Y)-phenylalanine -leucine YFL) being identical from yeast to human (**Figure 1A**). Notably, the α-carboxyl group of the carboxyl-terminal leucine is modified by methyl-esterification, also referred to as carboxy-methylation or methylation [17,18]. We refer to this modification as methylation, as is done by convention in the field. Because of the complex nature of the subunit nomenclature, we use gene names to refer to specific subunits throughout. This reversible modification is catalyzed by Leucine carboxyl methyltransferase 1 (LCMT1), which utilizes S-adenosyl-methionine (SAM) as the methyl donor [17,19–22]. For PP2Acα/β and PP4c it was shown that the modification is removed by protein phosphatase methylesterase 1 (PME1) [23–25]. In addition to hydrolyzing the methylester, PME1 binding reduces PP2A activity through a rearrangement of the PP2Acα active site and displacement of the two divalent cations, which are required for catalysis of the dephosphorylation reaction [26]. In cells, it is estimated that 70-90% of PP2Acα/β is methylated [18,27,28]. Similarly, prior reports indicate that approximately 75% of PP4c and PP6c are methylated in mouse embryonic fibroblasts [21].

**Figure 1.**
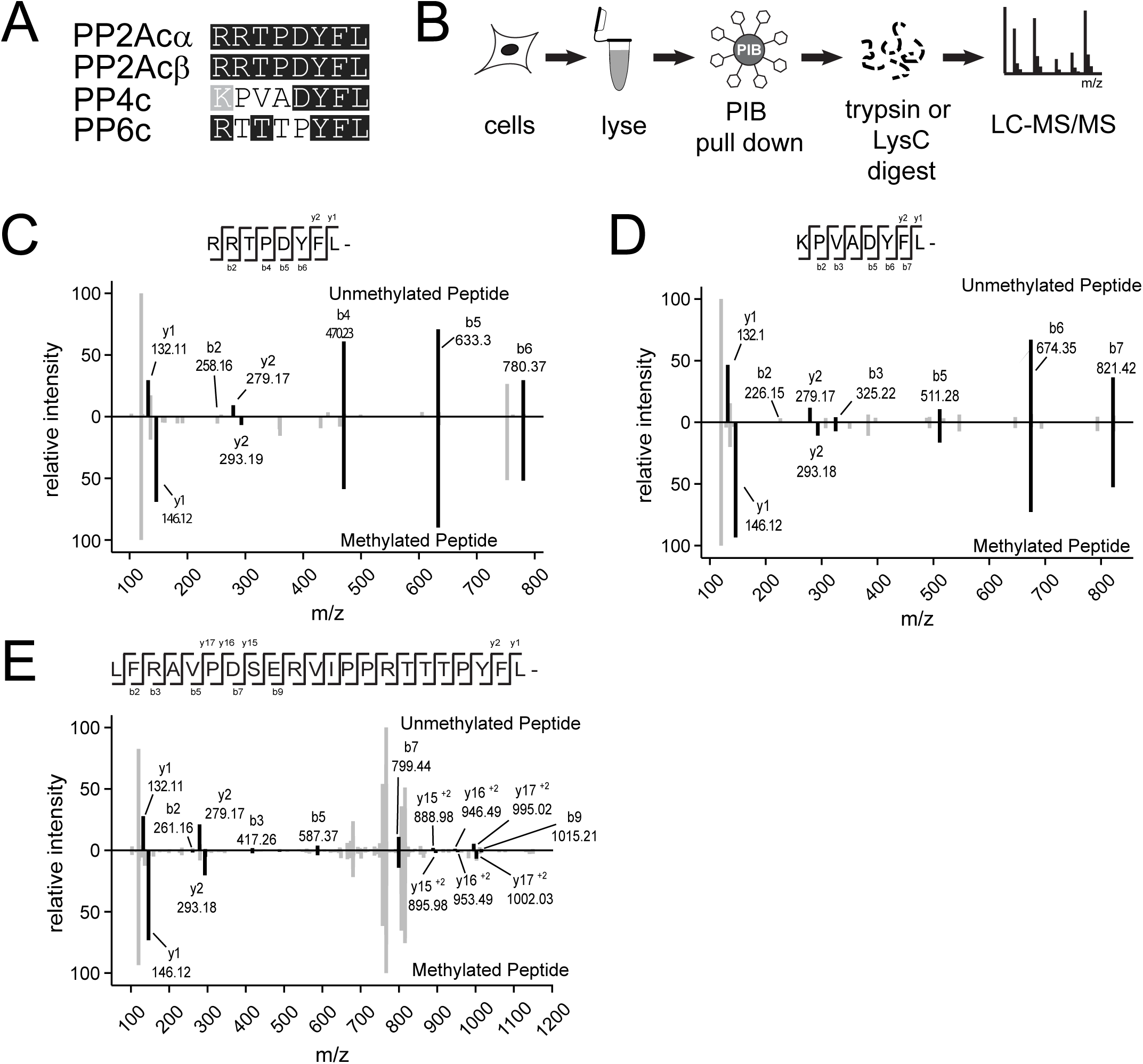
MS2 ion spectra of methylated and unmethylated C-terminal peptides for PP2Acα/β,, PP4c, and PP6.c. (A) Alignment of the last eight amino acids of the catalytic subunits of the PP2A, PP4, and PP6. (B) Phosphatase inhibitor bead () pulldown workflow. Cells were lysed. lysates and incubated with PIBs, elutes were digested overnight with either trypsin or LysC and analyzed using LC-MS/MS. (C-E) MS2 ion spectra of the C-terminal peptides for PP2Acα/β, B, PP4c, and PP6c. The unmethylated peptides are shown on the top half of each plot, and the methylated peptide is inverted on the bottom. Ions corresponding to the indicated peptide are shown in black, others are shown in grey.

The role of C-terminal methylation is best understood for PP2A. Methylation removes the negative charge of the carboxyl-terminus, thereby facilitating its docking into an acidic groove between the A and B subunits [5]. The deletion of the C-terminal leucine (ΔL309 mutation) of PP2A is frequently used as a mimetic of the unmethylated form. This mutation reduces the interaction of PP2Acα with B55, B56, and B72 subunits, while binding of the striatin family is increased [16,27]. Interestingly, viral antigens, such as the polyomavirus small T (ST) and middle T (MT) antigen, and the SV40 small tumor antigen (SVST) bind to the same extent to wild-type and ΔL309 PP2Acα/β [27,29]. Reduction of C-terminal methylation showed that B55 subunits preferentially bind to methylated PP2Acα/β, but that B56 and PR72/PR130 can be part of trimeric holoenzyme containing methylated or unmethylated PP2Acα/β [16,30]. For PP4c, loss of methylation reduces PP4R1 binding, while PP6 holoenzyme components bind PP6c independent of methylation [21].

These results demonstrate the important role of C-terminal methylation in PP2A and PP4 holoenzyme assembly. They qualitatively establish the ability of regulatory subunits to bind to either methylated or unmethylated PP2Acα/β. However, a quantitative comparison of the effects of the lack of the C-terminal leucine or methylation on subunit binding is missing. Many studies rely on epitope-tagging and exogenous expression because specific and well-characterized antibodies are not available for all subunit isoforms. Furthermore, several pan-PP2Acα/β antibodies have recently been shown to be biased and selectively recognize subpopulations of PP2Acα/β requiring a close re-examination and potentially reinterpretation of results from their use [31]. Thus, unbiased and quantitative analyses are warranted to evaluate the role of methylation in holoenzyme assembly and to establish if the deletion of the C-terminal leucine faithfully recapitulates the loss of methylation.

Here, we use mass spectrometry-based proteomics approaches combined with affinity enrichments, methylation-ablating mutations, and genome editing to comprehensively and quantitatively determine the role of C-terminal methylation in PP2A, PP4, and PP6 holoenzyme assembly. We directly compare all subunit binding to PP2Acβ, PP4c, and PP6c lacking the C-terminal leucine with changes in endogenous holoenzyme assembly upon disruption of LCMT1 expression. Using mass spectrometry, we overcome limitations of antibody-based subunit detection to investigate all subunits comprehensively. We generalize the role of C-terminal methylation in holoenzymes assembly across different transformed or non-transformed cell lines.

## Results

### Identification of C-terminal methylation of PP2Acα/β, PP4c, and PP6c by LC-MS/MS

We previously described a chemical proteomics approach to enrich, identify, and quantify endogenous PPPs from cells and tissues that we refer to as Phosphatase Inhibitor Beads combined with Mass Spectrometry (PIB-MS) [32]. Using a specific but non-selective PPP inhibitor microcystin-LR (MCLR) immobilized to sepharose beads, we can reproducibly enrich, identify, and quantify PP1α/β/γ, PP2Acα/β, PP4c, PP5c, and PP6c catalytic subunits as well as their associated scaffolding and regulatory subunits. Furthermore, PIB-MS enables the identification of PPP-associated PTMs [33]. To investigate C-terminal methylation of endogenous PP2Acα/β, PP4c, PP6c subunits, we enriched PPPs from the human cancer cell lines SW1088, C32, U-87, MCF-7, CAL-51 using PIBs, digested the isolated proteins into peptides, and analyzed them by liquid-chromatography coupled with tandem mass spectrometry (LC-MS/MS) (**Figure 1B**). We identified PP2Acα/β, PP4c, and PP6c carboxyl-terminal peptides modified by methyl-esterification in several of the cancer cell lines by database searching for a mass addition of 14.01565 Da (**Figure 1C-E, Supp. Table 1**).

### Methylation-ablating mutation affects holoenzyme assembly

To investigate the regulatory function of C-terminal methylation in PP2A holoenzyme assembly, PP2Acα/β mutants lacking the terminal amino acid leucine “ΔL” mutants have been employed [16,27,29]. We extended this approach to PP4c and PP6c by generating mammalian expression constructs in which the conserved C-terminal leucine was similarly removed (**Figure 1A**). Due to a lack of specific and well-characterized antibodies for all PP2A, PP4, and PP6 subunits, we chose a mass spectrometry-based proteomics approach. We comprehensively and quantitatively profile regulatory and scaffolding subunit interactions for the wild-type (WT) and ΔL catalytic subunits (**Figure 2A**). To distinguish between different isoforms of the same subunit family, all analyses were restricted to peptides that uniquely matched to the specific protein. We transfected FreeStyle 293F cells with 3xFLAG-tagged WT or ΔL mutant versions of PP2Acα/β, PP4c, or PP6c, affinity-enriched the catalytic subunits, and analyzed interacting proteins by label-free quantitative LC-MS/MS (**Figure 2A, Supp. Figure 1-3**). Using this approach, we reproducibly detected most of the regulatory and scaffolding subunits for PP2A and all subunits for PP4 and PP6 (**Supp. Table 2**). Furthermore, we quantified changes in scaffolding and regulatory subunit binding to the catalytic subunit in the presence and absence of the terminal leucine. These analyses revealed that the binding of the scaffolding subunits of PP2A, PR65α, and PR65β, was reduced by approximately 4-fold and 2-fold, respectively (**Figure 2B, Supp. Table 2**). Consistent with the literature [16,27], binding of the B55 family of PP2A regulatory subunits was reduced to less than 5% in the ΔL mutant compared to WT PP2Acβ. While the overall binding of B56 regulatory subunits was reduced, we also identified isoform-specific differences in regulatory subunit binding. B56α and B56ε were reduced by 32-fold and 11-fold, respectively, while B56γ and B56δ were reduced by 2-fold in binding to the ΔL mutant compared to WT PP2Acβ. This difference in subunit binding is consistent with unique amino acid sequence motifs in the C-terminal tail that distinguishes the B56α/β/ε versus B56γ/δ isoforms [34]. We also observed a strong reduction in the interaction of the ΔL catalytic subunit compared to WT for the B72 family. Finally, as previously shown, the members of the striatin family of regulatory subunits bound equally well to WT and ΔL mutant catalytic subunit. For PP4, we observed a reduction in PP4c-ΔL catalytic subunit interaction for PP4R1 and PP4R3α/β, with PP4R1 displaying the largest change in binding compared to WT (**Figure 2C**). The dependency of PP4R1 binding on PP4c methylation status was previously reported in mouse embryonic fibroblasts lacking LCMT1, while the binding of PPP4R3β was slightly, but not significantly reduced under these conditions [21]. We did not see any significant changes in subunit binding to PP6c-ΔL compared to WT (**Figure 2D**). Taken together, these data indicate that loss of the C-terminal leucine differentially affects subunit binding for each PP2A, PP4, and PP6. While for PP2Acα/β, loss of the C-terminal leucine strongly reduces the binding of most of the regulatory subunits, the same mutation had little effect on PP6c subunit interactions.

**Figure 2.**
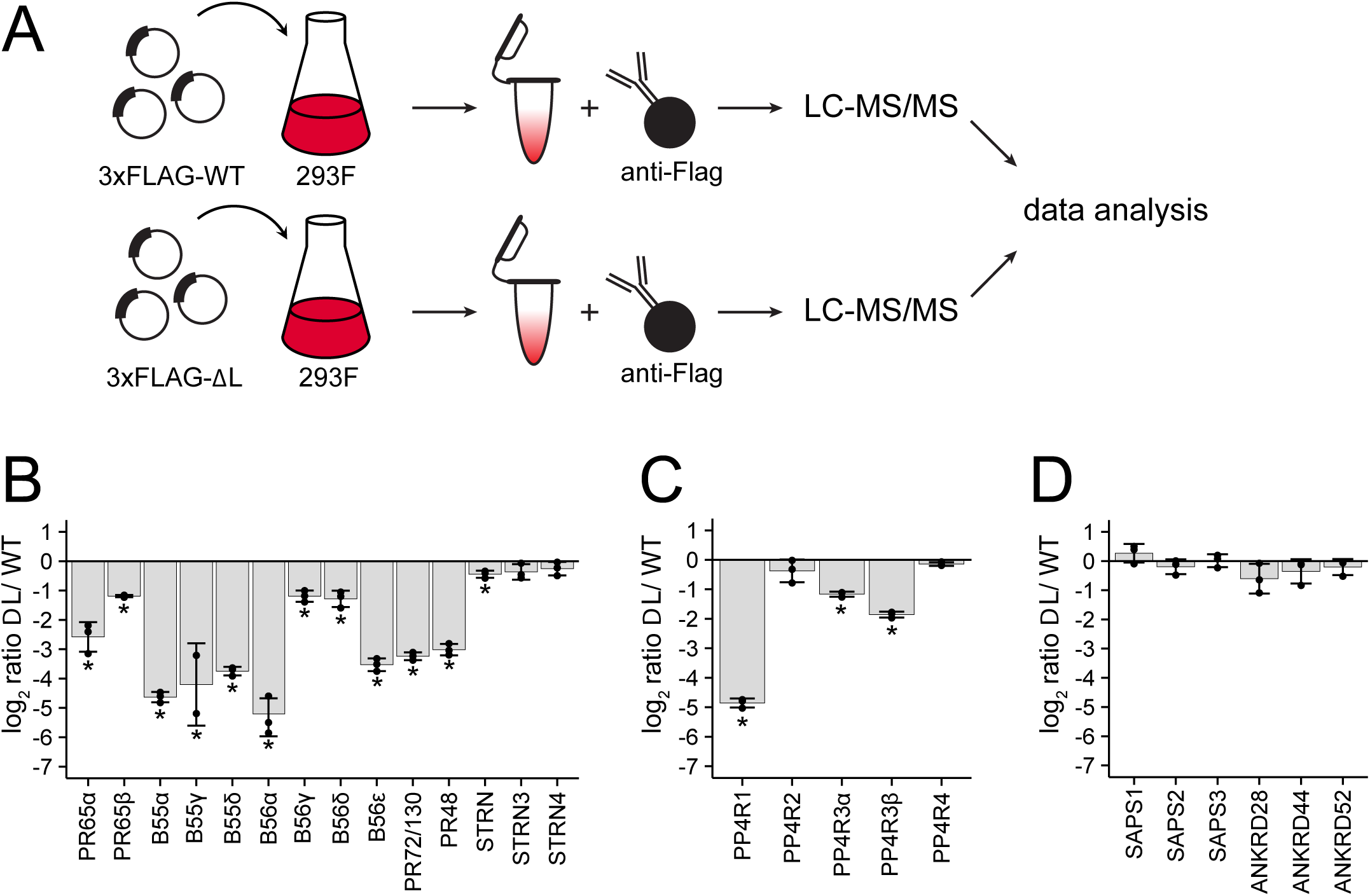
The Effects of deleting the C-terminal leucine on holoenzyme subunit composition of PP2A, PP4, and PP6. (A) Workflow for Flag affinity-purification mass spectrometry. Flag IPs were performed for PP2Acβ (B), PP4c (C), and PP6c (D) using either the wild-type or ΔL mutant and analyzed, and the log_2_-ratios are plotted. Individual ratios are represented. Error bars represent the standard deviation of three replicates. *, p < 0.05..

### Effects of LCMT1 depletion on PP2A, PP4, and PP6 subunit stability

The differences in holoenzyme assembly observed in the comparison of WT and ΔL mutant catalytic subunits could be due to a lack of methylation or the lack of the C-terminal leucine. To distinguish between these possibilities, we used CRISPR-Cas9 genome editing to disrupt LCMT1 in HEK293T cells [35,36]. We reasoned that because LCMT1 is an essential gene, LCMT1 knockout in mice results in embryonic lethality [28,37], we would likely select for hypomorphic knockouts in which most but not all alleles would be inactivated in triploid HEK293T cells. Therefore, we refer to these cells as a hypomorphic (HM). Briefly, HEK293T cells were infected with lentiviruses encoding Cas9 and a guide RNA targeting either exon 1 or 2 of LCMT1. After selection, single-cell clones were isolated and investigated for LCMT1 expression and PP2Acα/β methylation by Western blotting (**Figure 3A**). We found that LCMT1 abundance and PP2Acα/β methylation were reduced to nearly undetectable levels (**Figure 3A, B**). Taken together, these results suggest that most or all alleles of LCMT1 are disrupted, resulting in a decrease in PP2Acα/β C-terminal methylation.

**Figure 3.**
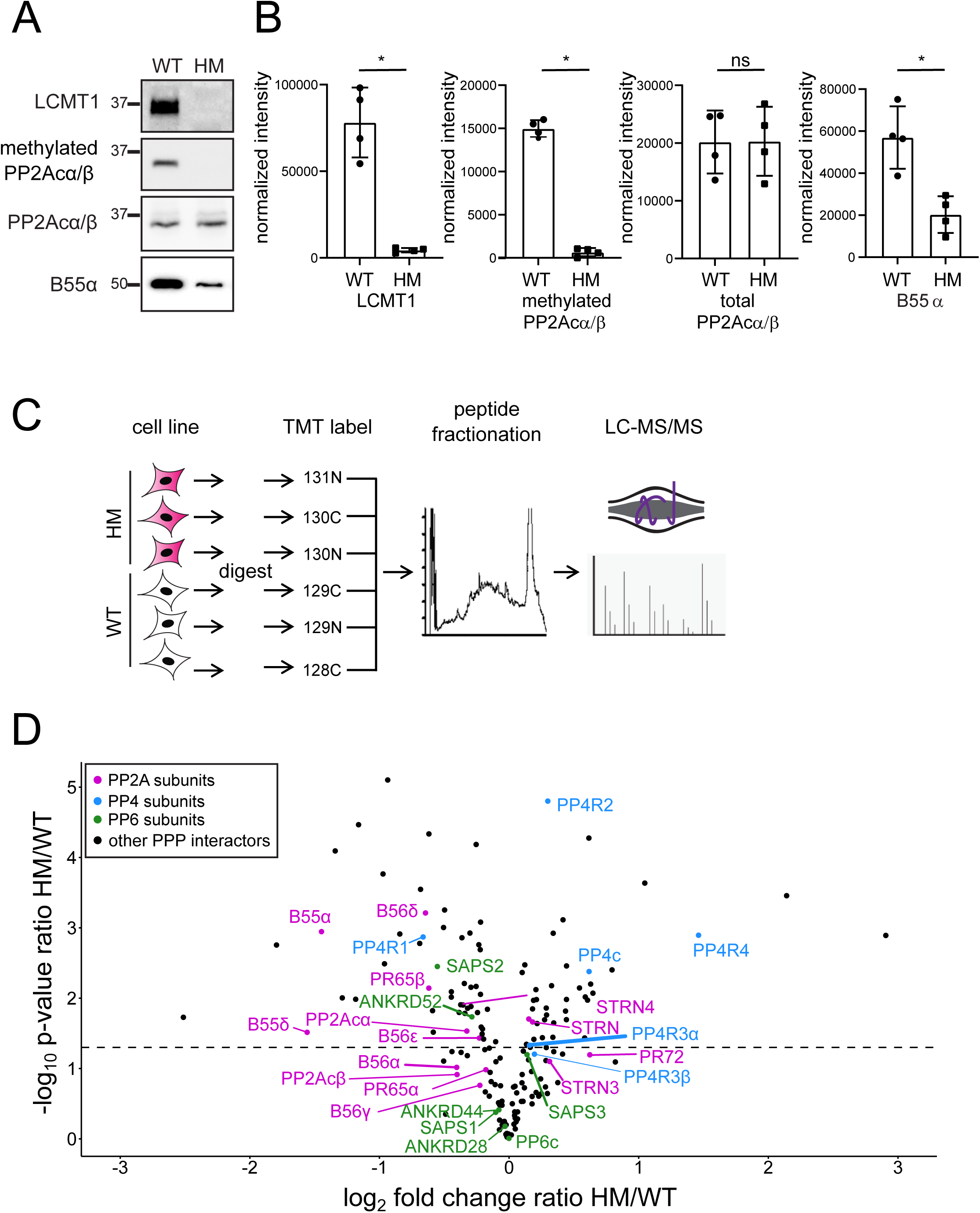
Disruption of LCMT1 in HEK293T cells. HEK293T cells were infected with lentiviruses encoding a gRNA targeting a region within LCMT1 exons 1 or 2. Cells were selected, and single colony screened for LCMT1 and methylated PP2A. (A) Wild-type (WT) and hypomorph (HM) lysates probed for LCMT1, methylated PP2Acα/β, total PP2Acα/β, and B55α. (B) Western blots were quantified, and individual data points are shown for LCMT1, methylated PP2Acα/β, total PP2Acα/β, and B55α intensities normalized to total protein loading. Error bars represent the standard deviation of four replicates. Mann-Whitney test was performed *, p < 0.05, ns – not significant. (C) Workflow for proteomic analysis of HEK293T LCMT1 HM cells compared to WTcells. (D) Volcano graph of known PPP interactors. The log_2_ fold change ratio HM/WT is plotted against the negative log_10_ of the p-value of the ratio HM/WT. A p-value cut-off of 0.05 (-log_10_ of 1.3) is indicated. PP2A subunits are labeled in purple, PP4 subunits are in blue, PP6 subunits are in green, and other PPP interactors are depicted as black dots.

B55 regulatory subunits preferentially bind to methylated PP2Acα/β and are degraded when unbound [16,21]. Consistently, we found that the abundance of the B55 subunit PPP2R2A was decreased upon LCMT1 reduction (**Figure 3B**). To comprehensively characterize total cellular protein abundance changes of PPP subunits upon LCMT1 disruption, we performed global quantitative proteomic analyses. Biological triplicates of wild-type (WT) and LCMT1 knockout cells were lysed, reduced, alkylated, and proteins were digested with trypsin into peptides. To quantitatively compare protein abundances across all samples, peptides were labeled with isobaric tandem mass tags (TMT), separated by off-line liquid chromatography to reduce sample complexity, and analyzed by LC-MS/MS (**Figure 3C, Supp. Table 3**). We found that all subunits of the B55 family were reduced in HM compared to WT cells (**Figure 3D**). Furthermore, the protein levels of B56 subunits were reduced in LCMT1 HM cells, albeit to a much lesser degree than B55 subunits (**Figure 3D**). Regulatory subunits belonging to the B72 or striatin family were, in most cases, not significantly changed in abundance (**Supp. Table 3**). We also found that PP4R1 was decreased by ∼2-fold, while the PP4R4 subunit was increased by ∼3-fold in HM compared to WT cells (**Figure 3D**). Finally, for PP6, only the SAPS2 subunit was slightly but decreased in HM cells. Regarding the catalytic subunits, we found that PP2Acα was decreased by ∼25%, while PP4c was increased by ∼50% in LCMT1 HM cells, suggesting that methylation stabilizes PP2Acα, but destabilizes PP4c. PP6c was unchanged between the two cell lines.

### PP2A, PP4, and PP6 holoenzyme assembly in LCMT1-depleted HEK293T cells

Next, we determined the effects of decreased C-terminal methylation on PP2A, PP4, and PP6 holoenzyme assembly. To do so, we used PIBs to enrich PPPs from HM and WT HEK293T cells. Consistent with our Western blot results (**Figure 3A, B**), our mass spectrometry analysis revealed that the C-terminal methylation of PP2Acα/β was reduced in HM compared to WT cells (**Figure 4A**). Similarly, PP4c, and PP6c C-terminal methylation was strongly decreased. The proteomic analyses revealed a decrease in PP2Acα abundance (**Figure 3D**), which was recapitulated in the PIB pulldowns (**Figure 4B**). The increase in PP4c abundance (**Figure 3D**) was also observed in the PIB analysis, while PP6c was mostly unchanged (**Figure 4B**).

**Figure 4.**
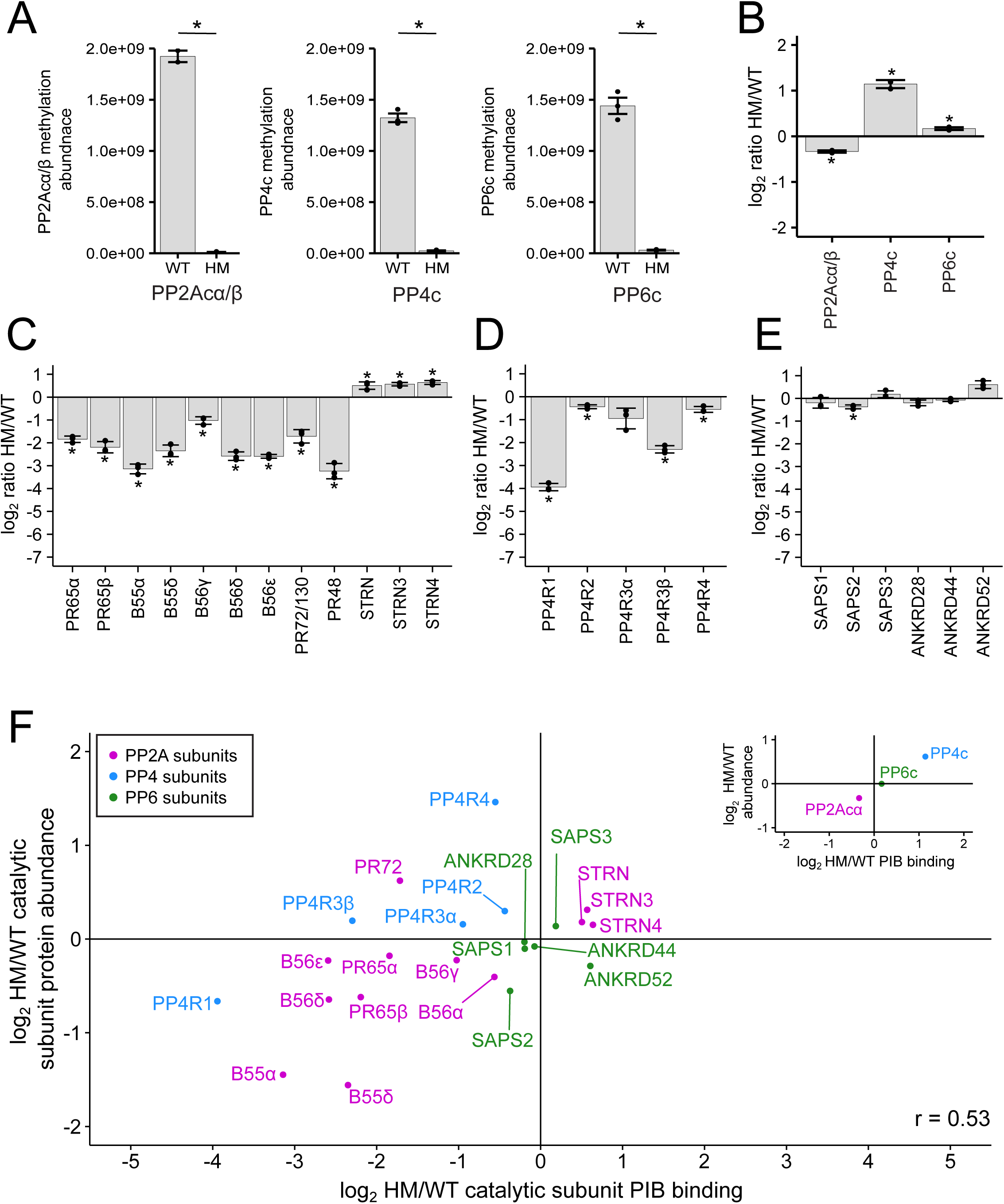
The effects of disrupting LCMT1expression in HEK293T cells on the holoenzyme composition of PP2A, PP4, and PP6. (A) Normalized peptides intensities of methylated peptides in wild-type (WT) and hypomorph (HM) cells by Western blot. Methylated peptides were normalized to the total protein amount of the respective catalytic subunit. Triplicates are plotted and shown as separate points. Error bars represent the standard deviation. The log_2_ ratio HM/WT of the protein abundance of the catalytic subunits (B), PP2A subunits (C), PP4 subunits (D), and PP6 subunits (E) enriched by phosphatase inhibitor beads (PIBs) and determined by label-free mass spectrometry. (F) Scatterplot depicting the log2 HM/WT ratio of PP2A, PP4, and PP6 subunits bound to PIBs and normalized to the respective catalytic subunit on the x-axis and the log_2_ HM/WT ratio protein abundance on the y-axis. PP2A subunits are labeled in purple, PP4 subunits are in blue, and PP6 subunits are in green. The scatterplot in the upper right depicts catalytic subunits PIB binding versus protein abundance. Two-sided Student’s t-test was performed, assuming unequal variance *, p < 0.05.

To identify changes in scaffolding and regulatory subunit binding, we normalized their abundances to the respective catalytic subunit (**Supp. Table 4**). For PP2A, we observed similar decreases in scaffolding and regulatory subunit binding upon LCMT1 disruption, as observed with the PP2Acβ -ΔL mutant. We found that B55, B56, and B72 subunits were decreased in binding to PP2Acβ, while members of the striatin family were increased upon decreased C-terminal methylation (**Figure 4C**). Similarly, for PP4, the subunit with the strongest decrease in binding to PP4c upon depletion of LCMT1 was PP4R1, while the lack of methylation reduced the PP4R3β interaction, but to a lesser degree (**Figure 4D**). PP4R2, PP4R3α, and PP4R4 were mostly unaffected by the reduction in methylation. Finally, for PP6, we did not detect any large changes in binding to PP6c (**Figure 4E**). Thus, in general, we observed the same trends in subunit interaction upon reduction of C-terminal methylation of the catalytic subunits as with the ΔL mutants, indicating that the ΔL mutation is a reasonable mimetic for the lack of methylation.

It is well established that B55 regulatory subunits are degraded when not incorporated into the PP2A holoenzyme complex, and that this incorporation is methylation-dependent [16,21]. To determine if the lack of holoenzyme incorporation affects the protein stability of other PP2A, PP4, and PP6 subunits, we compared the protein abundance and PIB binding data from HM and WT cells (**Figure 4F**). For the catalytic subunits, PIB binding correlated with protein abundance. In addition to B55 subunits, we found that PP4R1 and PP6R2 were decreased slightly in both PIB binding and protein abundances. This suggests that lack of incorporation might also lead to a decrease in protein stability of these free subunits. In contrast, although the B72 subunit PR72/PR130 was decreased in PP2Acα/β interaction by PIB-MS, it was increased in total cellular abundance. However, the increase in PP4R4 abundance in the HM cells did not lead to increased incorporation into PP4 holoenzymes. Taken together, these analyses indicate that certain subunits of PP2A and PP4 decrease in cellular protein abundance when not bound to a catalytic subunit, while others appear stable when not in phosphatase complexes.

### Generalizing the effects of reduced C-terminal methylation on holoenzyme assembly

To determine if the observed changes in holoenzyme assembly are unique to HM HEK293T cells or a general feature of LCMT1 disruption and reduction of C-terminal methylation, we generated LCMT1 deficient HeLa cells. HeLa cells were infected with lentiviruses encoding Cas9 and a gRNA targeting either exon 1 or 2 of LCMT1. After selection and single-cell cloning, LCMT1 and PP2Acα/β expression and PP2Acα/β C-terminal methylation were evaluated by Western blotting (**Figure 5A, B**). As shown for HEK293T cells by Western blotting, HeLa cells displayed a reduction in LCMT1 abundance and PP2Acα/β methylation, the latter of which was confirmed by mass spectrometry for all three catalytic subunits. While we readily observed the methylation of the C-terminus of PP2Acα/β, PP4c, and PP6c in wild-type HeLa cells, little to no modified C-terminal peptides were observed upon LCMT1 disruption (**Supp. Figure 4A**). To study the effects of C-terminal methylation on holoenzyme assembly in WT and HM HeLa cells, we performed PPP interactome analyses using PIBs (**Supp. Table 5**). We found that PP2Acα/β, PP4c, and PP6c mainly were unchanged in PIB binding (**Figure 5C**). For PP2A, we found that the interaction with the scaffolding subunit PR65cα/β was reduced (**Figure 5D**). We only identified and quantified two of the four isoforms of the B55 subunits (B55α and B55β), both of which were reduced by approximately 11- and 4-fold, respectively. For B56 subunits, we detected three isoforms (B56γ, B56δ, and B56ε). Similar to our analysis of the PP2Acβ-ΔL interaction, we observed that the B56ε isoform was 8-fold more reduced in binding to the catalytic subunit than B56γ and B56δ, supporting the notion of the differential affinity of the isoform groups for methylated PP2A catalytic subunit. We also found that PR48 was reduced, while striatins were mostly unchanged. The binding behavior of the PP4 and PP6 subunits recapitulated what we observed in HEK293T cells (**Figure 5 E, F**). In addition, we performed correlation analysis to directly compare the PP2A, PP4, and PP6 catalytic subunits interactions in HEK293T and HeLa cells, which yielded a high degree of concurrence between both datasets (Pearson correlation R = 0.85) (**Figure 5 G**).

**Figure 5.**
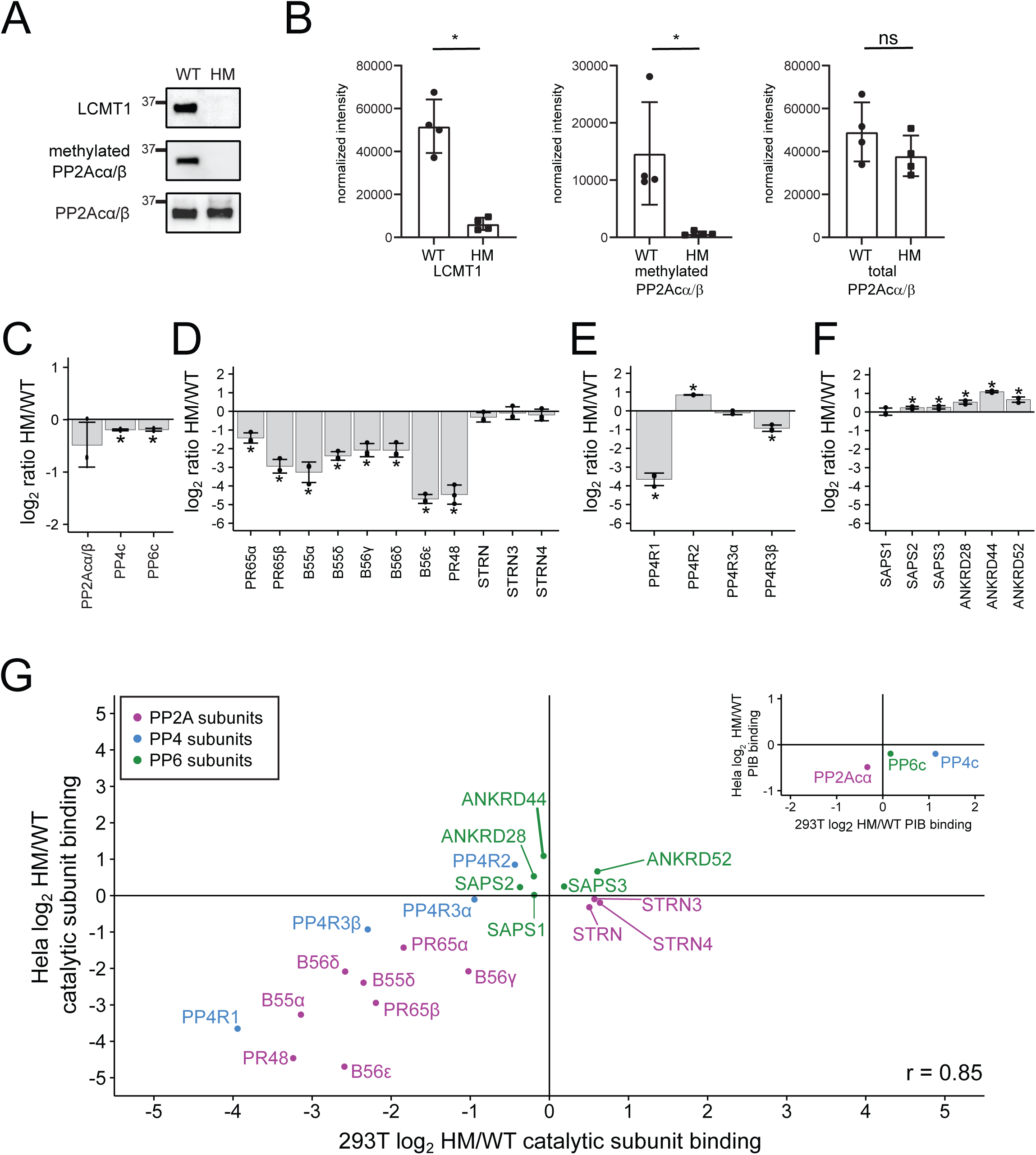
The effects of disrupting LCMT1 expression in HeLa cells on the holoenzyme composition of PP2A, PP4, and PP6. (A) Western blots wild-type (WT) HeLa and a hypomorph (HM) Hela clone probed for LCMT1, methylated PP2Acα/β, and total PP2Acα/β. (B) Western blots were quantified, and individual data points are shown for LCMT1, methylated PP2Acα/β, and total PP2Acα/β intensities normalized to total protein loading. Error bars represent the standard deviation of four replicates. Mann-Whitney test was performed *, p < 0.05, ns – not significant. The log_2_ HM/WT ratio of the abundance of the catalytic subunits (C), PP2A subunits (D), PP4 subunits (E), and PP6 subunits (F) bound to PIBs. (G) Scatterplot of the log_2_ HM/WT ratio of PP2A, PP4, and PP6 subunits PIB binding and normalized to the respective catalytic subunit in HEK293T versus Hela cells. PP2A subunits are labeled in purple, PP4 subunits are in blue, and PP6 subunits are in green. The scatterplot in the upper right depicts catalytic subunit PIB binding in both cell types. *, p < 0.05.

To investigate the role of C-terminal methylation in a non-transformed cell line, we disrupted LCMT1 expression in an hTERT-immortalized retinal pigment epithelial cell line (Rpe1). As described for HEK293T and HeLa cells, Rpe1 cells were infected with lentiviruses encoding Cas9 and a gRNA targeting either exon 1 or 2 of LCMT1. We have previously experienced problems with single-cell cloning of RPE1 and were unable to retrieve individual clones. Therefore, we tested the cell pool after selection to determine the abundance of LCMT1 and PP2Acα/β and the level of C-terminal methylation of PP2Acα/β by Western blotting (**Figure 6A, B**). This revealed a reduction of LCMT1 and methylated PP2Acα/β. The reduction of C-terminal methylation was confirmed by mass spectrometry. We found that the C-terminal methylation of PP2Acα/β, PP4c, and PP6c catalytic subunits was reduced to 11% or less (**Supp. Figure 4B**). Taken together, these data suggest that C-terminal methylation is sufficiently reduced in pooled LCMT1 HM Rpe1 cells to evaluate PP2A, PP4, and PP6 holoenzyme assembly by PIB-MS (**Supp. Table 6**). We observed an increase in PP2Acα binding to PIBs, which we currently cannot explain (**Figure 6C**). While the binding of the scaffolding, B55, B56, and B72 regulatory subunits to the catalytic subunit followed the trends observed in HEK293T and HeLa cells (**Figure 6C**), striatins did not. In other cell lines and employing the ΔL mutant in FreeStyle 293F cells, we observed that the interaction of striatin with the catalytic subunit was methylation independent. Indeed, we observed a slight increase or no change in striatin binding upon LCMT1 disruption in HEK293T and HeLa cells, respectively (**Figure 4C, 5D**). However, in Rpe1 cells, Striatin binding was decreased upon reduction of C-terminal methylation by 1.5-3-fold, depending on the specific subunit (**Figure 6D**).

**Figure 6.**
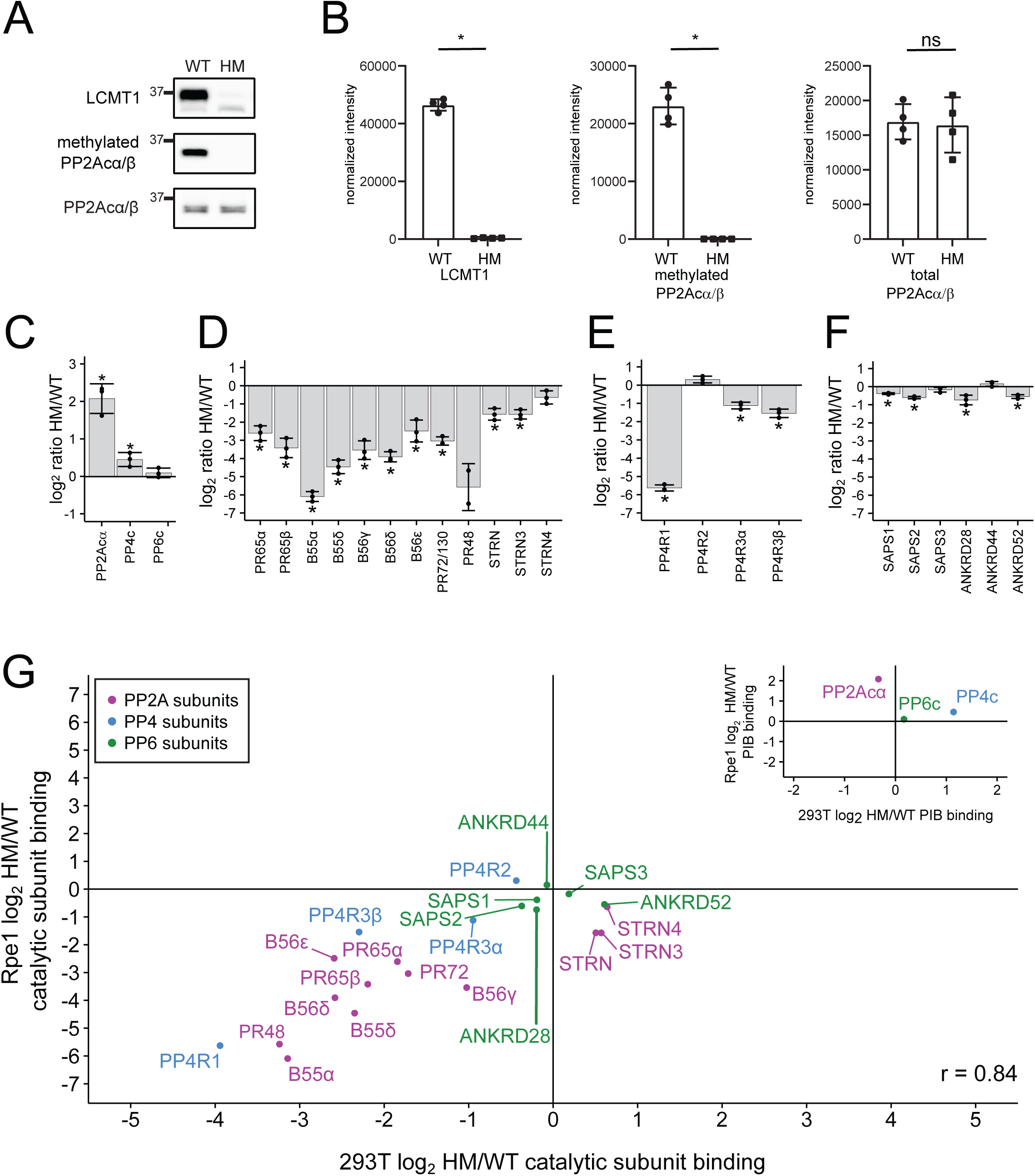
The effects of disrupting LCMT1 expression in Rpe1 cells on the holoenzyme composition of PP2A, PP4, and PP6. (A) Western blots of wild-type (WT) and hypomorph (HM) Rpe1 cells probed for LCMT1, methylated PP2Acα/β, and total PP2Acα/β (B) Western blots were quantified and correct for loading based on total protein, and individual data points are shown for LCMT1, methylated PP2Acα/β, and total PP2Acα/β. Error bars represent the standard deviation of four replicates. The log_2_ HM/WT ratio of the abundance of the catalytic subunits (C), PP2A subunits (D), PP4 subunits (E), and PP6 subunits (F) bound to the PIBs as determined by proteomics analysis. (G) Scatterplot of the log_2_ HM/WT ratio of PP2A, PP4, and PP6 subunits binding to PIBs and normalized to the respective catalytic subunit from 293T cells versus Rpe1 cells. PP2A subunits are labeled in purple, PP4 subunits are in blue, and PP6 subunits are in green. The scatterplot in the upper right depicts catalytic subunit PIB binding 293T cells versus Rpe1 cells. *, p < 0.05.

For PP4 and PP6, the results were consistent with our observation in HeLa and HEK293T cells (**Figure 6 E, F**). The general consistency of the observed trends, besides striatins, is also observed in the correlation analysis of methylation-dependent changes in catalytic subunit interactions upon LCMT1 disruption in HEK293T and Rpe1 cells (R = 0.84) (**Figure 6G**).

## Discussion

PPPs are responsible for the majority of cellular dephosphorylation and regulate most biological processes [1]. PPPs achieve substrate specificity by forming multimeric holoenzyme complexes with various regulatory subunits. The plethora of PPP subunits expands the number of potential enzymes from 10 catalytic subunits to hundreds of holoenzymes. For PP2A, PP4, and PP6, holoenzyme formation is at least in part controlled by C-terminal-methylation [16,21,27,38–40]. Determining how C-terminal methylation regulates PP2A, PP4, and PP6 holoenzyme composition is important because changes in holoenzyme assembly are linked to human diseases, including Alzheimer’s Disease (AD) and cancer. For instance, PP2A-B55α, is the main phosphatase that dephosphorylates tau [41]. Tau hyperphosphorylation is associated with the formation of neurofibrillary tangles, a hallmark of AD [42–44]. Reduced PP2Acα/β, methylation has been associated with tau hyperphosphorylation in AD brains [45,46], and LCMT1 is downregulated in AD neurons with neurofibrillary tangles [46].

Because of the large number of subunits, investigations of the endogenous protein using specific antibodies is challenging. Thus, previous studies have often relied on epitope-tagging, exogenous expression, and detection of PP2A catalytic subunit binding using antibodies. Recently, it was shown that several PP2Acα/β antibodies preferentially recognize the unmethylated forms or cross-react with PP4c and PP6c, resulting in detection bias [31,47,48]. Monitoring or immunoprecipitating PP2Acα/β using these antibodies provides an incomplete, biased picture of the role of C-terminal methylation, requiring a close re-examination and potentially reinterpretation of results from their use. Furthermore, prior comparisons of subunit binding were mostly qualitative, establishing if a specific subunit family and their various isoforms can bind methylated, unmethylated, or ΔL mutant forms of PP2A, PP4, and PP6 catalytic subunits. Therefore, an unbiased, quantitative mass spectrometry-based analysis of the impact of C-terminal methylation on holoenzyme assembly is warranted.

We used two different approaches to investigate the effects C-terminal methylation has on the holoenzyme assembly of the PP2A, PP4, and PP6. First, we exogenously expressed either the wild-type (WT) or a C-terminal leucine deleted mutant (ΔL mutant) of each catalytic subunit in Freestyle 293F cells, an approach commonly used to mimic loss of methylation [16,27]. We immunoprecipitated the WT and ΔL catalytic subunits and quantified subunit interactions by mass spectrometry. Second, we disrupted LCMT1 expression in three different cell types (HEK293T, HeLa, and Rpe1) and investigated the holoenzyme assembly of endogenous catalytic subunits by activity-based enrichment using PIB-MS [32]. Using both approaches, we observed a reduction of B55, B56, and B72 subunit binding to PP2Acα/β. Thus, our analyses demonstrate that the ΔL mutation is a valid approach to investigate the effect of C-terminal methylation of catalytic subunits of the PP2A, PP4, and PP6.

For the B55 and striatin families of PP2A regulatory subunits in 293T and HeLa cells, our results are consistent with previous reports demonstrating that B55 subunits strongly rely on C-terminal methylation for holoenzyme formation, while striatins do not [16,27,37,38,40,49,50]. Recently, we found that the catalytic subunit of PP2A is phosphorylated at the C-terminus at Thr304 during mitosis and that this phosphorylation also disrupts B55 subunit binding and could affect methylation occupancy [33]. However, the stoichiometry of this phosphorylation is low and would only have a minor effect on methylation [51]. In Rpe1 cells, we observed a 4-fold reduction in binding of STRN and STRN3 upon decreasing C-terminal methylation. Rpe1 cells are immortalized but not transformed, which could contribute to the observed behavior. We are currently exploring if this differential rewiring of PP2A-Striatin holoenzyme complexes is a general feature distinguishing transformed from non-transformed immortalized cells.

In contrast, the role of C-terminal methylation in B56 holoenzyme assembly is less clearly defined. In a qualitative analysis, B56 binding to the PP2A catalytic subunit was shown to be methylation-independent [16]. Both methylated and unmethylated PP2Acα/β are present in B56 holoenzyme complexes [16]. In bovine brain extract, no unmethylated B56-containing trimeric holoenzymes could be detected, and B56 regulatory subunits displayed a lower affinity for unmethylated compared to methylated PP2Acα/β [39]. In *S.cerevisiae*, binding of the B56 homolog was strongly reduced upon loss of methylation [52]. Structural analysis of a B56γ containing PP2A holoenzyme revealed that methylation promotes holoenzyme formation but is not required *in vitro* [5]. Our results suggest that while B56 subunits can interact with methylated and unmethylated PP2Acα/β, methylation promotes this association. For the ΔL mutation, we observed a difference in the degree of the reduction in binding for different B56 isoforms. While PPP2R5A and E were reduced by 32-fold and 11-fold, respectively, B56γ and B56δ were reduced by only 2-fold in binding. However, only LCMT1 disruption in HeLa cells resulted in a similar distinction in B56 isoform interactions with PP2Acα/β, suggesting that this might be a cell type-specific effect.

Little is known about the methylation-dependency of the PR48 and PR72/130. B72 isoforms were shown to interact with the ΔL mutant, methylated, and unmethylated PP2A catalytic subunit [16]. However, inhibition of PME1 increased the association of PR72/130 with the PP2A catalytic subunit, suggesting that it preferentially associates with methylated PP2Acα/β [30]. We find that binding of PR48 and PR72/130 to the ΔL mutant and to the PP2A catalytic subunit upon disruption of LCMT1 expression is reduced. Thus, while B72 subunits can interact with unmethylated PP2Acα/β, our data suggest that B72 isoforms strongly prefer binding to methylated PP2Acα/β.

For PP4 and PP6, the analyses of catalytic subunit binding upon deletion of the C-terminal leucine and disruption of LCMT1 expression revealed the largest fold-change in catalytic subunit interaction for PP4R1, while PP4R3α/β were modestly reduced. Finally, we did not observe large changes in PP6 subunit binding, indicating that PP6 holoenzyme assembly is mostly methylation-independent.

Also, some subunits are unstable when not incorporated into a holoenzyme complex [16,37,49]. Consistent with previous reports, we found that B55 subunit abundances were reduced upon LCMT1 disruption and lack of holoenzyme incorporation. We also observed that B56 subunits and PPP4R1 were slightly decreased. Intriguingly, PPP4R4 was increased in abundance in HEK293T cells upon LCMT1 depletion. Thus, while protein abundance changes of PPP subunits can correlate with holoenzyme formation, direct determination of holoenzyme composition is needed to determine the specific effects.

The comparable trends in methylation-dependency of holoenzyme assembly using the ΔL mutant and deletion of LCMT1 in three different cell types suggest a conserved regulatory relationship between C-terminal methylation and specific subunit binding. In all systems investigated, we saw a strong reduction in LCMT1 levels and C-terminal methylation. Likely, modulation of LCMT1 levels will differentially affect subunits depending on the stoichiometry of C-terminal methylation and subunit affinity for the methylated C-terminus. Indeed, it was previously shown that the B55 family had a higher affinity for the methylated C-terminus of PP2Acα/β than B56 [16,27,40]. Further investigations of the relationship of C-terminal methylation stoichiometry and PP2A regulatory subunit binding in cells are needed to elucidate these dependencies. In most cell types and tissues [18,21,27,28,40], methylation stoichiometry is high, establishing a repertoire of holoenzymes responsible for cellular dephosphorylation. Not only are B55 and B56 regulatory subunits differentially sensitive to lack of methylation, but they also display different substrate specificities, with B55 containing holoenzyme opposing proline-directed kinases and B56 containing holoenzymes opposing basophilic kinases [53,54]. Thus, changes in methylation and thereby the holoenzyme repertoire differentially affect kinase opposition. Future studies will need to determine which phosphorylation sites are specifically sensitive to these changes and identify the downstream signaling pathways that are differentially regulated.

## Materials and Methods

### Cell culture

HEK293T, Rpe1, and HeLa cells were grown as adherent cultures in Dulbecco’s modified Eagle’s medium (DMEM) (Corning) with 10% fetal bovine serum (FBS) (Hyclone) and penicillin-streptomycin (100 U/ml and 100ug/ml, respectively, Corning) at 37°C, 5% CO_2_. Freestyle 293F cells were grown in suspension using FreeStyle 293 media (Gibco), with 10% FBS at 37°C, 5% CO_2_. 293FT cells cultured in IMDM (Lonza) with 8% FBS and 1% pen/strep at 37°C, 5% CO_2_.

### LentiCrispr-Cas9; cloning, viral production, and infection

Creating the lentiCrispr virus was performed as previously described [35]. Briefly, guide RNAs were selected using CRISPOR and CHOPCHOP [55,56]. Oligonucleotides were synthesized by IDT. LentiCRISPRv2 was digested with BsmbI (NEB) for 1 hour followed by the addition of 1µl of CIP (NEB) and incubated at 37°C for 30 minutes. The backbone was gel purified. To anneal the gRNAs they were resuspended in 25 µl of nuclease-free water and 10 µl of each guide were combined with 5 µl of NEB buffer 3 and incubated at 95°C for 5 minutes, then immediately moved to a 70°C water bath for 10 minutes, at which point the heat was turned off and samples cooled in the bath overnight. The annealed oligonucleotides were phosphorylated by combining 5 µl of the annealed oligos, 12 µl water, and 2 µl of 10x Promega T4 ligase buffer (contains ATP) with 1µl of PNK kinase (NEB) and incubated at 37°C for 1 hour, then incubated at 95°C for 5 minutes. A ligation reaction was performed using 1µl of digested and CIP treated pLentiCRISPRv2, 5 µl of phosphorylated oligonucleotides, 7.5 µl of 2x Promega quick ligation buffer, and 1ul of Promega T4 polymerase, which incubated at room temperature (RT) for 30 minutes. After, 5 µl of the ligation product was transformed into competent DH5α. Plasmids were verified by sequencing. For creating the lentivirus, 293FT cells were transfected with LentiCRISPRv2-g1-4-LCMT1, pMD2.G, and psPAX2 at a 5:2:3 µg ratio using PEI. PEI was removed after 4 hours and fresh media without antibiotics was added. The media was collected and replaced after 48 hours and collected again after 96 hours. The media was centrifuged at 2000 rpm for 5 minutes then filtered through a 0.22µm PVDF membrane, aliquoted and frozen at -80°C. For infection, roughly 1 million cells were incubated for 48 hours with 1ml of lentivirus. After, the cells were selected against with puromycin at 1µg/ml for 1 week. Clones were collected and tested for the expression of LCMT1 and methylated PPP2CA/B by SDS-PAGE.

### PP2Acβ, PP4c, and PP6c mutagenesis and transfections

Human PP2Acβ, PP4c, and PP6c were cloned into p3XFLAG-CMV10 vector using HindIII and EcoRI restriction sites and verified by sequencing. Leu 309 in PPP2CB, Leu 307 in PP4c, and Leu 305 in PP6C were deleted by inserting a stop codon using Quick Change Lightning Kit (Agilent) as per manufacturer’s instructions. All mutations were verified by sequencing. The FreeStyle 293F cells were grown to a confluency of 1×10^6^ cells/mL and then transfected with 0.5 mg/L DNA and 1.5 mg/L polyethylenimine (PEI)/ with p3XFLAG-CMV10-PP2Acα/β B/PP4c/PP6c WT or ΔL. Briefly, 0.25 mg DNA was diluted in 25 mL Opti-MEM Reduced Serum media. Separately, 0.75mg of sterile PEI was diluted in 25 mL Opti-MEM Reduced Serum media. PEI solution was added to the DNA solution, inverted, and incubated for 20 min at room temperature. After incubation, DNA:PEI mixture was added to 500 mL of Expi293 cells. Cells were incubated with skaking at 125 rpm for 48 hours. Cells were collected and pelleted at 1500 rpm for 2 minutes and frozen at -80°C until the purification was performed.

### Western blotting, detection, and densitometric analysis

For Western blot analyses, cells were lysed in Laemmli samples buffer, separated by SDS-PAGE, and analyzed by western blotting. Antibodies were diluted in 3% milk in TBST. Antibodies used in the study: Total PP2A (11H12) and methylated PP2A (7C10) (gift from Dr. Egon Ogris), B55α (#sc-365282, Santa Cruz Biotechnology), LCMT1 (sc-362551, Santa Cruz Biotechnology), andFLAG (F3165, Sigma). Mouse and rabbit HRP-linked secondary antibodies were purchased from Jackson Immunoresearch. Western blots were developed using Clarity Western ECL substrate (BioRad) and visualized using a BioRad ChemiDoc station. Quantification was carried out in Fiji ImageJ[57].

### PIB and FLAG pulldowns

FLAG pulldowns were carried out as previously described [58]. Briefly, 293F cells expressing either PP2Acβ, PP4c, or PP6c WT or ΔL were collected and lysed in lysis buffer (50 mM Tris pH 7.5, 500 mM NaCl and 0.5% (w/v) Triton X-100, with an EDTA-free protease inhibitor tablet (Roche) per 10 mL of lysis buffer). FLAG-tagged proteins were purified using FLAG M2 affinity gel (Sigma) rotated end over end for 3 h at 4°C. The beads were washed five times with lysis buffer and eluted with 3XFLAG peptide (final concentration 150 ng µl^-1^). The elutes were aliquoted for western blot or TCA precipitation. The samples for western blot analysis were heated with 2X Laemelli buffer at 95°C for 5 minutes. The eluates for TCA precipitation were precipitated using 20% tri-chloroacetic acid (TCA)(Sigma), washed twice with 10% TCA and twice more with acetone (Burdick & Jackson, Muskegon, MI) and digested overnight in 25 mM ammonium bicarbonate (ambic) with 1:200 (w/w) dilution of trypsin (Promega) for mass spectrometric analysis. PIBs were generated and used as described before [32]. Elutes were processed as previously described for FLAG pulldowns.

### LC-MS/MS analysis

FLAG-pulldowns were analyzed on a Q-Exactive Plus quadrupole Orbitrap mass spectrometer (ThermoScientific) equipped with an Easy-nLC 1000 (ThermoScientific) and nanospray source (ThermoScientific). Peptides were resuspended in 5% methanol / 1% formic acid and loaded on to a trap column (1 cm length, 100 μm inner diameter, ReproSil, C18 AQ 5 μm 120 Å pore (Dr. Maisch, Ammerbuch, Germany)) vented to waste via a micro-tee and eluted across a fritless analytical resolving column (35 cm length, 100 μm inner diameter, ReproSil, C18 AQ 3 μm 120 Å pore) pulled in-house (Sutter P-2000, Sutter Instruments, San Francisco, CA) with a 45 minute gradient of 5-30% LC-MS buffer B (LC-MS buffer A: 0.0625% formic acid, 3% ACN; LC-MS buffer B: 0.0625% formic acid, 95% ACN). The Q-Exactive Plus was set to perform an Orbitrap MS1 scan (R=70K; AGC target = 1e6) from 350 – 1500 m/z, followed by HCD MS2 spectra on the 10 most abundant precursor ions detected by Orbitrap scanning (R=17.5K; AGC target = 1e5; max ion time = 50ms) before repeating the cycle. Precursor ions were isolated for HCD by quadrupole isolation at width = 1 m/z and HCD fragmentation at 26 normalized collision energy (NCE). Charge state 2, 3, and 4 ions were selected for MS2. Precursor ions were added to a dynamic exclusion list +/-20 ppm for 15 seconds. Raw data were searched using COMET (release version 2014.01) in high resolution mode [59] against a target-decoy (reversed) [60] version of the human proteome sequence database (UniProt; downloaded 2/2020, 40704 entries of forward and reverse protein sequences) with a precursor mass tolerance of +/-1 Da and a fragment ion mass tolerance of 0.02 Da, and requiring fully tryptic peptides (K, R; not preceding P) with up to three mis-cleavages. Static modifications included carbamidomethylcysteine and variable modifications included: oxidized methionine. Searches were filtered using orthogonal measures including mass measurement accuracy (+/-3 ppm), Xcorr for charges from +2 through +4, and dCn targeting a <1% FDR at the peptide level. Quantification of LC-MS/MS spectra was performed using BasicQuan or MassChroQ and the iBAQ method [61,62].

PIB-pulldowns were analyzed on a Lumos Fusion Orbitrap mass spectrometer (ThermoScientific) equipped with an Easy-nLC 1200 (ThermoScientific). Peptides were resuspended in 5% methanol / 1% formic acid across a column (45 cm length, 100 μm inner diameter, ReproSil, C18 AQ 1.8 μm 120 Å pore) pulled in-house across a 45 min gradient from 3% acetonitrile/0.0625% formic acid to 37% acetonitrile/0.0625% formic acid. The Orbitrap Lumos was set to perform an Orbitrap MS1 scan (R=120K; AGC target = 2e5) from 350 – 1500 m/z, followed by HCD MS2 spectra in top-speed mode for 1.5 sec by Orbitrap scanning (R=15K; AGC target = 2.5e4; max ion time = 50ms) before repeating the cycle. Precursor ions were isolated for HCD by quadrupole isolation at width = 1.1 m/z and HCD fragmentation at 28% HCD collision energy. Charge state 2, 3, and 4 ions were selected for MS2. Precursor ions were added to a dynamic exclusion list +/-20 ppm for 18 seconds. Raw data were searched using COMET (release version 2014.01) in high resolution mode [59] against a target-decoy (reversed) [60] version of the human proteome sequence database (UniProt; downloaded 2/2020, 40704 entries of forward and reverse protein sequences) with a precursor mass tolerance of +/-1 Da and a fragment ion mass tolerance of 0.02 Da. For tryptic digests, fully tryptic peptides were required (K, R; not preceding P) with up to three mis-cleavages. Static modifications included carbamidomethylcysteine and variable modifications included: oxidized methionine. For Lys-C digests, cleavage C-terminal to lysine was required with up to three mis-cleavages. Static modifications included carbamidomethylcysteine and variable modifications included: oxidized methionine. For the identification of C-terminal methylation +14.01565Da as a variable modification of the carboxyl-terminus was included. Searches were filtered using orthogonal measures including mass measurement accuracy (+/-3 ppm), Xcorr for charges from +2 through +4, and dCn targeting a <1% FDR at the peptide level. Quantification of LC-MS/MS spectra was performed using BasicQuan or MassChroQ and the iBAQ method [61,62]. For missing values, retention time alignment and smart quantification was performed.

### TMT quantitative proteomics

HEK293T cells were collected and lysed in lysis buffer (8 M urea and 50 mM Tris pH 8.1 with protease inhibitors) in the presence of protease inhibitors. After lysis, a small aliquot was removed for BCA (Pierce) analysis, and the remaining lysate was reduced with 5 mM DTT at 50°C for 30 min followed by alkylation with 15 mM iodoacetamide in dark for 1 h. The lysates were then diluted fivefold with 25 mM Tris pH 8.1 and digested overnight with trypsin (1:100 w/w) at 37°C. The digests were desalted using C_18_ solid-phase extraction cartridges (ThermoFischer Scientific) and an aliquot of each of the desalted eluates corresponding to 40 µg of peptide digest was dried by vacuum centrifugation in separate tubes. For TMT-labeling, acetonitrile was added to a final concentration of 20% in 166 mM HEPES, and peptides were transferred to dried, individual TMT 11-plex reagent (ThermoFisher Scientific), vortexed, and mixed with the reagent. After 1 h at room temperature, an aliquot was removed for label-check. Once labeling efficiency was confirmed to be above 95%, each digest was quenched with 5 µl of 500 mM ammonium bicarbonate solution for 10 min, combined, diluted threefold with 0.1% TFA in water, and desalted using C_18_ solid-phase extraction cartridges (ThermoFisher Scientific). The desalted multiplexes were dried by vacuum centrifugation, and separated on a pentafluorophenyl (PFP) analytical column [XSelect HSS PFP XP column, Waters; 100 A, 2.5 mm inner diameter (ID), 150 mm in length] into 48 fractions by gradient elution. The 48 fractions were reduced into 16 fractions and dried by vacuum centrifugation [63]. Fractions were analyzed using an Easy LC-1000 (Proxeon) and Orbitrap Fusion (Thermo Fisher Scientific) LC-MS/MS platform across a 2-hour gradient from 3% acetonitrile/0.0625% formic acid to 37% acetonitrile/0.0625% formic acid [64]. The Orbitrap Fusion was operated in data-dependent, SPS-MS3 quantification mode wherein an Orbitrap MS1 scan was taken (scan range = 350 – 1300 m/z, R = 120K, AGC target = 2.5e5, max ion injection time = 100ms)[65,66]. Followed by data-dependent Ion trap MS2 scans on the most abundant precursors for 3 seconds. Ion selection; charge state = 2: minimum intensity 2e5, precursor selection range 550-1300 m/z; charge state = 3-5: minimum intensity 3e5. Quadrupole isolation = 0.7 m/z, AGC target = 1e4, max ion injection time = 40ms, CID collision energy = 32%, ion trap scan rate: rapid. Orbitrap MS3 scans for quantification (R = 50K, AGC target = 5e4, max ion injection time = 86ms, HCD collision energy = 60%, scan range = 100 – 700 m/z, synchronous precursors selected = 10). The raw data files from Orbitrap Fusion were searched using COMET with a static mass of 229.162932 on peptide N-termini and lysines and 57.02146 Da on cysteines, and a variable mass of 15.99491 Da on methionines against the target-decoy version of the human proteome sequence database (UniProt; downloaded 2/2020, 40704 entries of forward and reverse protein sequences) and filtered to a <1% FDR at the peptide level. Quantification of LC-MS/MS spectra was performed using in house developed software [58].

### Data analysis and statistics

Data analysis was carried out using R statistical software (https://www.r-project.org/, version 4.0.2) in conjunction with R studio (version 1.2) [67] or GraphPad Prism. All plots were created using the ggplot2 package [68]. For proteins in the FLAG and PIB pulldowns to be considered for further analysis, they had to be identified with a total peptide count greater than 1 (TP > 1) and at least one unique match (UM ≥ 1). Calculations were based only on quantifications of unique matched peptides to distinguish isoforms. Protein abundances were log-transformed. For FLAG and PIB, protein log_2_ ratios were calculated by subtracting each replicate of the mutant condition (ΔL, HM) from the median of the WT. For Western blot quantifications. p-values were calculated by Mann-Whitney test. For proteomics analysis, p-values were calculated using a two-tailed Student’s t-test, assuming unequal variance.

## Acknowledgments

We thank Brooke Brauer and Dr. Isha Nasa, as well as other members of the Kettenbach and Gerber labs for helpful discussions and comments. We would also like to thank Dr. Egon Ogris for the PP2A (11H12) and methylated PP2A (7C10) antibodies. LentiCRISPR v2 was a gift from Feng Zhang (Addgene plasmid # 52961). This work is supported by grants R35GM119455 from NIH/NIGMS and R33CA225458 from NIH/NCI to ANK. The shared resources of Norris Cotton Cancer Center are supported by grant P30CA023108 from the NIH/NCI. The BioMT tissue culture core is supported by grant P20GM113132 from NIH/NIGMS.

## Author contributions

SPL and ANK designed experiments and wrote the manuscript. SPL, ECG, and LEC performed experiments. SPL analyzed data. MEA generate and supported the proteomics workflow.

## Conflicts of interest

The authors have no conflict of interest.

## Notes

### Competing Interest Statement

The authors have declared no competing interest.

