## Supplemental Figure 1 for "Regulation of PP2A, PP4, and PP6 holoenzyme assembly by carboxyl-terminal methylation"

Supp. Figure 1

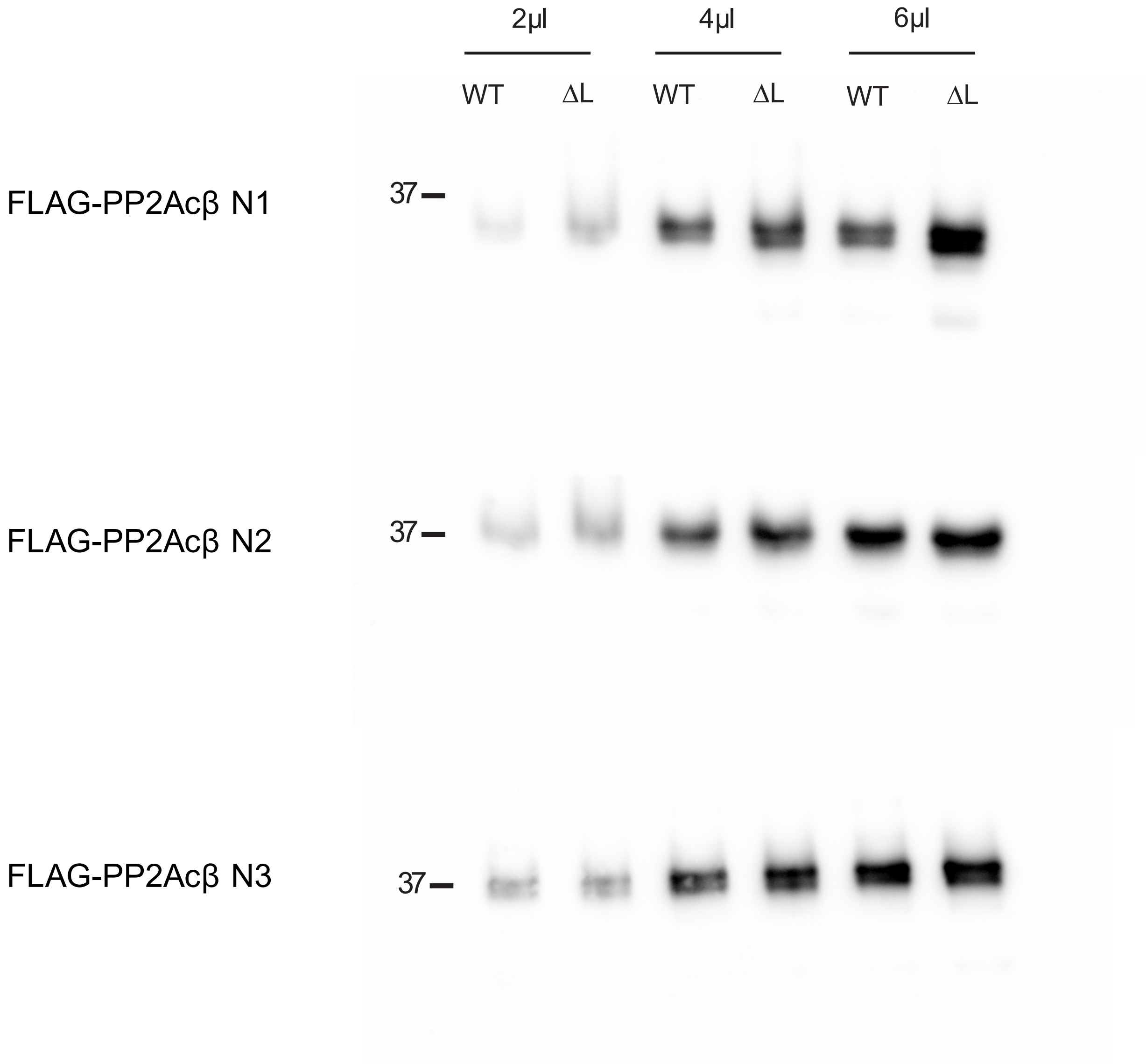


**Supp. Figure 1**. Purification of wild-type or ΔL mutant catalytic subunits of PP2A. Western blots of affinity purified FLAG-PP2Acβ wild-type and ΔL mutant.
