## Supplemental Figure 2 for "Regulation of PP2A, PP4, and PP6 holoenzyme assembly by carboxyl-terminal methylation"

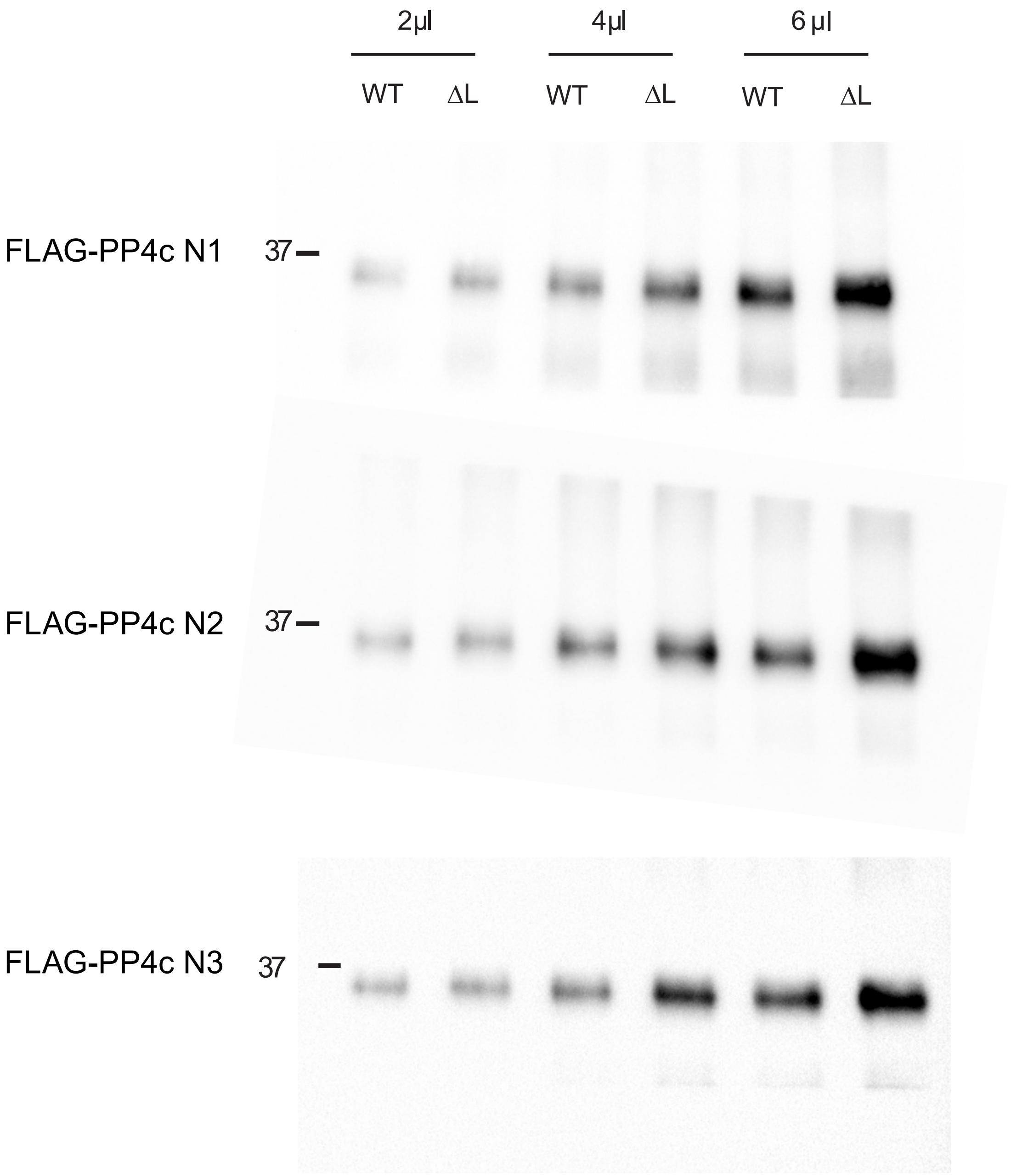


**Supp. Figure 2**. Purification of wild-type or ΔL mutant catalytic subunits of PP4. Western blots of affinity purified FLAG-PP4c wild-type and ΔL mutant.
