## Supplemental Figure 3 for "Regulation of PP2A, PP4, and PP6 holoenzyme assembly by carboxyl-terminal methylation"

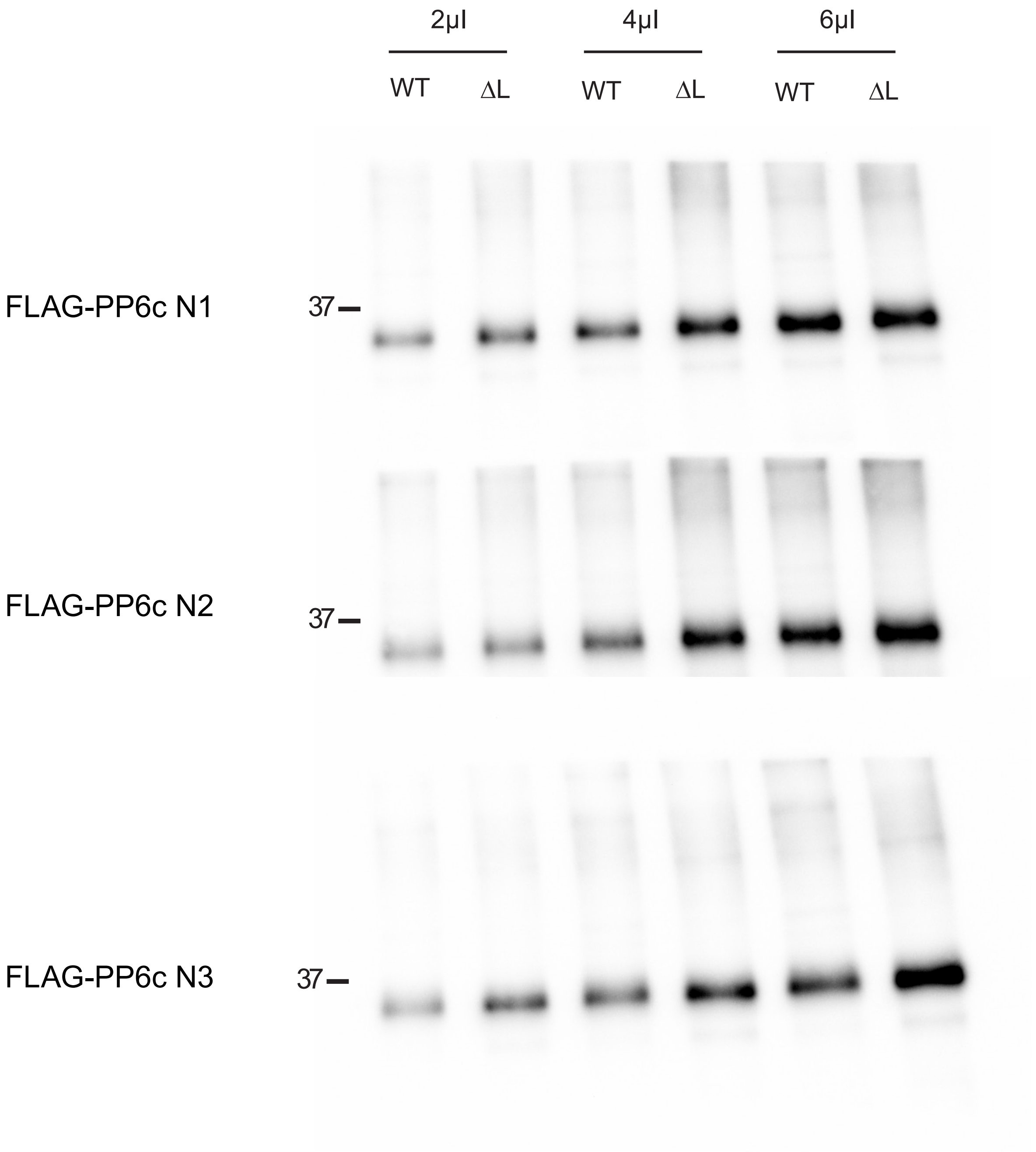


**Supp. Figure 3**. Purification of wild-type or ΔL mutant catalytic subunits of PP6. Western blots of affinity purified FLAG-PP6c wild-type and ΔL mutant.
