## Supplemental Figure 4 for "Regulation of PP2A, PP4, and PP6 holoenzyme assembly by carboxyl-terminal methylation"

Supp. Figure 4

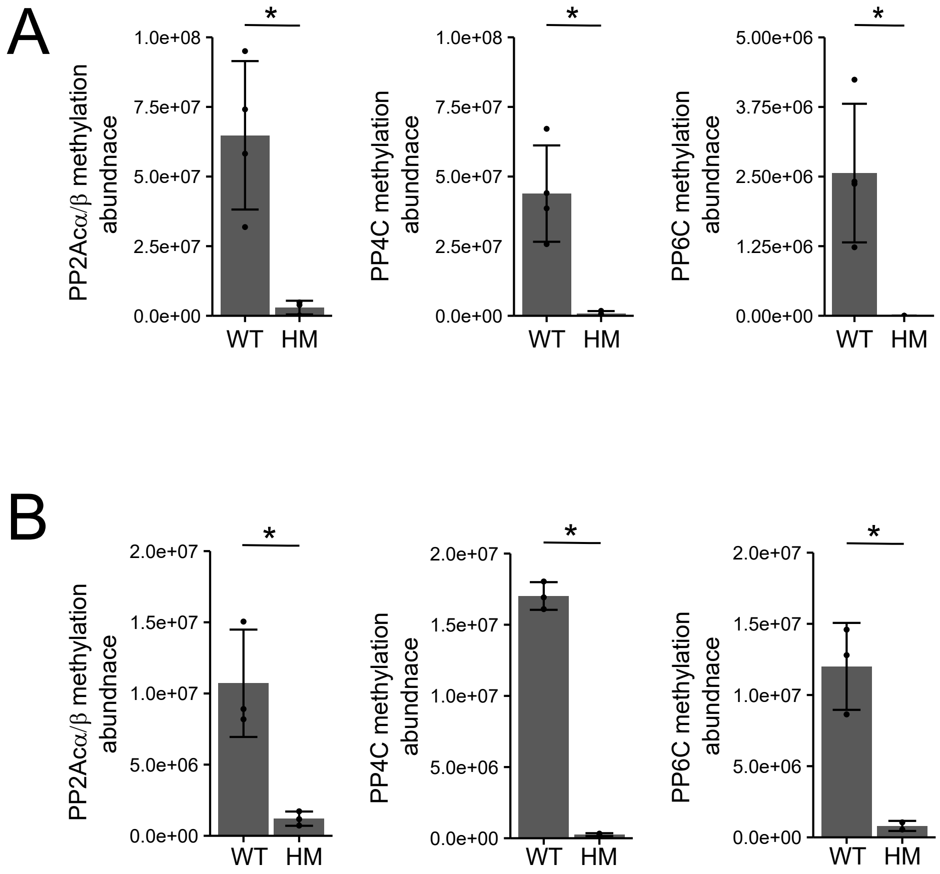


**Supp. Figure 4**. Quantification of C-terminal methylation using mass spectrometry of LCMT1 disrupted HeLa and Rpe1 cells. Normalized peptides intensities of methylated peptides in WT and HM in Hela (A) and Rpe1 (B) cells. Methylated peptides were normalized to the amount of the respected catalytic subunit. Triplicates are plotted and shown as separate points. Error bars represent the standard deviation. The $\log_2$ ratio HM/WT of the abundance of the catalytic subunits. Two-sided student's t-test was performed, assuming unequal variance, * $p < 0.05$.
